# Bayesian multilevel modeling of group and participant level effect sizes: Stronger inter-individual differences in pupil dilation compared to EEG alpha power during aversive conditioning

**DOI:** 10.1101/2025.11.07.687228

**Authors:** Sarah M. Gardy, Jourdan J. Pouliot, Faith E. Gilbert, Richard T. Ward, Caitlin Traiser, Payton Chiasson, Andreas Keil, Andrew Farkas

**Author notes:** Corresponding Author: Sarah M. Gardy, Department of Psychology, Laboratory for Brain, Body, and Behavior, University of Florida.

## Abstract

Aversive conditioning prompts reliable changes in the power of EEG alpha-band oscillations and pupil dilation. Both variables have been used to test hypotheses on the acquisition, generalization, and extinction of conditioned threat. Existing studies have largely relied on trial averages and group-level analyses. Thus, the variability of these physiological markers to aversive learning at the subject level is currently unknown. Comparisons of group-level analyses in prior studies suggest that pupil dilation and EEG-alpha activity capture complementary information. However, to date, no study has directly compared these two markers in terms of their effect sizes at the level of individual participants. The present study employed Bayesian multilevel modeling to quantify the variability of conditioning effect estimates for alpha-band power and pupillometry. Estimates were examined at the group level and at the participant level, across two conditioning paradigms, involving visual and auditory cues. Although the two metrics shared similar effect sizes at the group level, participant-level variability in these effect sizes was substantially higher for pupil-dilation compared to alpha-power, and this finding was replicated across both paradigms. These findings have important implications for clinical and inter-individual difference research which requires both the quantification of effects at the participant-level as well as meaningful variability between-participants that can be linked to relevant differences such as anxiety.

## 1 Introduction

Identifying and responding to cues that predict aversive outcomes is an essential adaptive behavior. Furthermore, the degree of responses to such cues is expected to differ across individuals and has been associated with maladaptive personality traits and disorders (Kaczkurkin et al., 2017; Kaczkurkin & Lissek, 2013; Lissek, 2012; Lissek et al., 2014). Learned aversive responses are readily studied in laboratory settings via Pavlovian conditioning paradigms in which a neutral cue (CS+) is paired with an aversive outcome (US) such as loud noise or noxious shock (Fullana et al., 2020; Lonsdorf et al., 2017). Through repeated pairings, the CS+ evokes defensive behavioral and physiological responses, which reflect aversive learning. Differential aversive conditioning extends this paradigm by measuring the response to an unpaired neutral stimulus (CS-).

It has been well established that various physiological measures reflect aversive conditioning over a group of participants (Lonsdorf et al., 2017). These measures include pupil dilation and EEG-derived alpha-band power, both of which reliably show differential responses to CS+ vs. CS-stimuli (Friedl & Keil, 2020a; Panitz et al., 2019a; Pietrock et al., 2019). Pupillary responses are highly sensitive to a range of cognitive and physical factors, including arousal, task demand, ambient light, and the proximity or salience of stimuli (Bradley et al., 2008, 2017; Steinhauer et al., 2022). This includes work linking pupil dilation to changes in autonomic states during cognitive, affective, and conditioning tasks (Friedl & Keil, 2020a). For example, research comparing the anticipation of an electric shock to neutral acoustic stimuli found that anticipation of the aversive shock resulted in increased pupil dilation compared to neutral trials (Bitsios et al., 2004). Pupil dilation offers enhanced sensitivity to changes in arousal over other physiological markers such as skin conductance and heart rate variability (Korn et al., 2017; Lonsdorf et al., 2017; Partala & Surakka, 2003).

At the level of human neurophysiology, selective transient power changes in the EEG alpha band (8-13 Hz) have emerged as another robust metric accompanying aversive conditioning. Transient alpha suppression is thought to reflect neural disinhibition in sensory regions mediated by thalamic and cortical areas (Li & Keil, 2023; for review see Klimesch, 2012). Exposure to CS+ stimuli has been shown to prompt such decreases in alpha-band power over parietal cortices, when using visual (Friedl & Keil, 2020b, 2021; Panitz et al., 2019b) and auditory cues (Farkas et al., 2024). For instance, Friedl and Keil (2021) found selectively reduced parieto-occipital alpha power when observers viewed grating stimuli at aversively conditioned spatial positions, compared to unconditioned positions.

Although the evidence for group-level effects of aversive conditioning on alpha-power and pupil-dilation has been extensive, what is unclear is how these measures compare at the level of individual participants and across different conditioning paradigms. There is strong interest in understanding inter-individual differences in aversive conditioning, given its relevance for theories of anxiety and post-traumatic stress (Beckers et al., 2023; Craske et al., 2008; Seligowski et al., 2019). Quantifying the effect sizes and variability of each metric at the level of individual participants would enable researchers to assess and interpret inter-individual differences as meaningful. This relates to a growing concern, known as the reliability paradox, that some paradigms produce consistent group-level effects because there are near equivalent effects per person (Hedge et al., 2018). For example, it is possible that a physiological measure and paradigm could produce near identical results among participants during conditioning and there would be no variation based on anxiety. To reach the goal of translating psychophysiological findings for more precise transdiagnostic information (Cuthbert & Kozak, 2013; Kotov et al., 2017), research using multimodal recording and more sophisticated statistics are needed to better identify the paradigms and measures that provide meaningful individual differences.

The present study utilizes two datasets (Farkas et al., 2024; Pouliot et al., 2025) to compare simultaneously recorded alpha and pupil across different conditioning paradigms. The studies differ in the sensory modality of the cues; one paired a noxious noise with a neutral tone CS+ while the other paired a noxious noise with a face CS+. This provides the ability to examine whether modality of conditioned stimuli influences the effect sizes of alpha and pupil measures. Central to the aims of the study, we examined if and how the effect varied between participants via Bayesian multilevel models. This allowed for the modeling of the nested structure of trials and participants within study to obtain regularized and reliable participant-level estimates (Gelman et al., 2012). To more easily compare the CS+ vs CS-effects per person across modalities and measures, a single-trial effect size estimate of the CS+ vs CS-contrast was obtained by dividing the mean conditioning effect by the estimated noise per trial. This effect size estimate is similar to the equation of the well-established Cohen’s D (Cohen, 2013), but it is more directly interpretable because it represents the difference between the conditions scaled by noise at the single-trial level. By contrast, a standard *group-level* Cohen’s D would change as a function of the number of trials averaged. The present effect size estimates also benefit from the multilevel structure of the hierarchical model, which provides regularization of the mean differences and estimates of single-trial error, making the contrast estimates more robust. Thus, this approach allows us to directly compare group-level and participant-level effect size posteriors for alpha power changes and pupil diameter, across the two paradigms.

## 2 Methods

### 2.1 Participants

This study comprised two experiments with separate samples, one utilizing tone stimuli and another using face stimuli. A total of 136 participants were included in the present analysis. Of these, 135 provided usable EEG alpha power data, with an average age of 20.63 (18-47 age range). The breakdown by sex included 101 females and 34 males, with 97 participants identifying as women, 34 as men, and 4 as non-binary. The sample included 9 Asian, 6 Black, 113 White, 6 other, and 1 who preferred not to answer. Additionally, 27 individuals identified as Hispanic. For the pupil data, 101 participants were included (mean age: 20.22 years; range 18-47), of whom 78 were female. In the sample, 75 identified as women, 23 as male, and 3 as non-binary. The racial breakdown included 6 Asian, 6 Black, 86 White, and 3 identifying as other.

Participants were required to be at least 18 years of age, have normal to corrected normal vision, and have no history of epilepsy. All participants gave written consent to participate in the study. Compensation varied by study: participants in the tone study received either course credit or $20 per hour and those in the face experiment received either course credit or a $70 remuneration. Both experiments adhered to the Declaration of Helsinki and received approval from the Institutional Review Board at the University of Florida.

### 2.2 Tone Stimuli and Paradigm

In the auditory experiment, participants were presented with an aversive conditioning task that paired an aversive noise with a soft tone CS+. Before the experiment, participants underwent a prescreening process for Misophonia. Following this, participants completed demographic questionnaires along with assessments related to Misophonia, social anxiety, and depression. The questionnaires are not examined in the current study.

During the experiment, participants were exposed to three tones of different pitch (low, medium, and high), presented in a randomized order across multiple trials, each lasting four seconds. Stimuli were presented using Psychtoolbox (Brainard, 1997) on a Cambridge Research Systems Display ++ monitor (1,920 × 1080 pixels, 120 Hz refresh rate), positioned approximately 120 cm in front of the participant. Participants sat approximately 60 cm from the eye tracker lens. The tone stimuli comprised three sinewave tones, each lasting four seconds and sampled at 22,000 Hz. To prevent abrupt onset and offset sound spikes, a cosine square window with a 20-point ramp-on and ramp-off was applied at the beginning and end of each tone. The three pitch conditions – CS+, GS1, and GS2 (here labeled the CS-) – were determined by an exponential function that modulated the sinewaves at 320 Hz, 541 Hz or 914 Hz, respectively. During the Acquisition phase, the 91 dBA white noise served as the unconditioned stimulus (US) and co-terminated with the CS+ tone during the final second of its presentation. To facilitate learning, the first and third trials of the acquisition phase were always CS+ trials, while the sequence of all remaining trials was pseudorandomized to ensure no more than two CS+ trials occurred consecutively.

The tone aversive generalization task consisted of 240 trials divided into two phases habituation and acquisition, each containing 120 trials. In each trial, one of the three tones was played for four seconds while participants fixated on a white dot (0.5° visual angle) displayed on the monitor. Trials were separated by an inter-trial interval (ITI) ranging from 1.85 to 3.5 seconds. One specific tone (CS+) was consistently paired with a loud noise (91 dBA white noise) during the final second of its presentation in 80 trials.

### 2.3 Face Stimuli & Paradigm

In the face experiment, participants were presented with an aversive conditioning task that paired an aversive noise with a face. Prior to the task, participants answered questionnaires related to social anxiety. During the conditioning task, participants were seated approximately 60 cm from the eye tracker and 120 cm from the display monitor. There was a total of 291 trials during which facial stimuli were presented in the center of the screen at a visual angle of –7.63 degrees. During the trials, the US was paired with the CS+ at 100% reinforcement rate. The first 11 trials served as booster trials and exclusively presented the CS+, and two stimuli that were the most perceptually dissimilar to the CS+ averaged together (CS-). This was done to ensure participants’ familiarity with the unaltered stimuli before encountering generalization phase. Across the remaining 280 trials, all faces were presented in a pseudo-random order. The CS+ was presented for 3 seconds, with the US co-terminating during the last second. All other faces were presented for 2 seconds each. All faces were flickered at a 15Hz frequency with a central fixation dot presented during the inter-trial interval (2-4 seconds in duration).

Faces were manipulated across a feature gradient which involved seven monochrome female faces displayed against a dark gray background. Four of these faces were morphs generated using iMorph software. Each morph blended the central face (CS+) with one of the other original, perceptually dissimilar faces (CS-‘s). To ensure consistency in contrast, all facial stimuli were processed with a Gaussian filter (full-width half maximum of 50 grayscale units). The unconditioned stimuli (US) consisted of a burst of 92 dB white noise presented during the last second of the CS+ presentation, bilaterally from two speakers located behind the participant. The MATLAB Psychophysics Toolbox (Brainard, 1997) was used to program and execute the experiment.

### 2.4 Pupil Data Acquisition and Processing

Eye-tracking data were recorded with the EyeLink 1000 Plus system, using a 16 mm lens positioned in front of the monitor. The system operated at a sampling rate of 500 Hz. Pupil diameter was determined for each participant individually by fitting an ellipse to the pupil mass threshold. The system’s infrared illumination level was initially set to 100% and modified as needed based on each participant’s pupil size and corneal reflection. Calibration and validation were conducted using a nine-point grid, where a white circle (1° visual angle) was sequentially displayed at each of the nine locations against a black background. If the system failed to detect the participant’s pupil diameter at any point, adjustments were made to the lens position and pupil thresholds until stable tracking was achieved.

Missing segments of pupil data were interpolated using a cubic spline method to ensure continuity in the signal. All preprocessing code is available on the OSF repository https://osf.io/ebs2a/overview. For each participant, average pupil dilation responses were computed across trials. Analysis focused on the final 500 ms of the CS presentation before US onset. To control for baseline differences, pupil data were baseline-adjusted by subtracting the average of 200 ms pre-trial period from each trial. Lastly, pupil size measurements were converted from their original units to millimeters, following best practices (Steinhauer et al., 2022).

### 2.5 EEG Data Acquisition and Processing

EEG data were recorded at 500 Hz sampling rate with a 128-channel HydroCel net. Impedences were kept below 60 kΩ. For both experiments, continuous EEG data were referenced to the grand-averaged reference. Signal processing occurred in MATLAB (version 2024, Natick, MA, USA) with a version of the Statistical Correction of Artifacts in Dense Array Studies (SCADS) method (Junghöfer et al., 2000). In both experiments, the signal was filtered using a 3rd order Butterworth low-pass filter with a 3dB point at 40 Hz and a 3^rd^ order high-pass filter with a 3 dB point at .1 Hz. In the tone experiment, data were segmented into 2600 ms trial epochs, spanning 600 ms pre-tone onset to 2000 ms post-tone onset. In the face experiment, data were segmented into 3000 ms trial epochs, spanning 800 ms pre-face onset to 2200 ms post-face onsets.

Segmented trials were assessed for artifact contamination using SCADS, which evaluates mean amplitude, standard deviation, and gradient voltage amplitude (Junghöfer et al., 2000). Channels were marked as globally bad if artifact-related metrics exceeded 2.5 standard deviations above the median quality index. These globally bad channels were interpolated with 2D spline interpolation. at the trial level, a quality index threshold of 2.5 standard deviations was applied, where trials exceeding this threshold were flagged and corrected via within-trial 2D spline interpolation. Eye artifacts were corrected, and rejection was conducted with a regression-based EOG method (Schloegl et al., 2009; Schlögl et al., 2007).

Alpha-band activity was quantified using Morlet wavelet analysis conducted in MATLAB (version 2024, Natick, MA, USA). Artifact-free single trials were transformed into the time-frequency domain using a family of Morlet wavelets with a fixed parameter (m = 10). Wavelet convolution was applied to the data across frequencies ranging from 1.15 and 35 Hz, in steps of 3.08 Hz. These settings resulted in a frequency uncertainty (sigma_f_) of 1.04 Hz and a time uncertainty (sigma_t_) of 153 ms at the chosen frequency of 10.38 Hz. An average of 8.64 channels were interpolated per participant and an average of 49.33 trials removed across the tone study. For the face stimuli, an average of 8.97 channels were interpolated per participant and an average of 62.93 trials were removed per participant.

### 2.6 Bayesian Models and Analyses

This study utilizes two Bayesian models for statistical analyses. According to recommended practice, Bayesian models were built incrementally with parameters added to each subsequent model until the final models were reached (McElreath, 2020). The multilevel structure specified in the models allowed for reliable granular estimates through a by-default regularization (Gelman et al., 2012). In the present study, this allowed for the estimation of credible conditioning effects and effect sizes per participant via regularization toward the averages of the group. The models were specified in the Stan programming language (Stan Development Team, 2024) that estimates posteriors using a form of Hamiltonian Monte Carlo sampling method (Duane et al., 1987; Hoffman & Gelman, 2011). In line with recommended practices, priors were selected to be either uninformative (wide relative to the scale of the data) or to represent the multilevel structure of the data (Gelman et al., 2014; McElreath, 2020). A total of 40,000 samples were taken to estimate the posteriors across 4 Markov Chains and 10,000 samples each. All chains converged based on the R-hat metric cutoff criteria at 1.02 for the estimated parameters.

The alpha and pupil models were designed to be as similar as possible to allow for comparisons of uncertainty between the measures. The main difference was the use of a log-normal likelihood function for alpha-power whereas a normal distribution was used for pupil-dilation. A log-normal distribution is justified for alpha-power because the measure is strictly positive, skewed, and is roughly normal after a log-transformation. Using the log-normal likelihood function is similar to transforming the data into decibels which is a routine preprocessing procedure in EEG spectral analysis (Delorme et al., 2011). The second difference was that alpha-power was not baselined per trial, as alpha power statistics are typically computed on non-adjusted data (Keil et al., 2022). Baseline models were designed and tested for alpha and fit much more poorly in terms of estimated leave-one out log-likelihood cross-validation accuracy; this suggests using the baseline led to overfitting and a poorer estimation of the underlying data generative process (McElreath, 2020, Chapter 7). The uninformative priors were adjusted to account for differences in physiological measurement between alpha (mv) and pupil (mm), informed by prior predictive analyses and previous literature. While this study only considers the most dissimilar cues to the CS+ which we label as CS-, mean effects for generalization stimuli that looked similar to the CS+ were also assessed to provide a better estimation of single-trial error (σ).

For Model 1, alpha amplitude for each observation comes from a log normal distribution, in which the mean (µ) and a standard deviation (σ) are on the log-scale. Each participant mean, µ, is determined by the additive association between participant’s intercept and the effect of cue based on condition. As is shown in the following equations, a posterior for each intercept (β0), cue (βCS), and single-trial error (σ) was found per participant. Each was regularized via a multilevel prior of the group-average posterior. All other priors were selected to be weakly informative based on prior predictive analyses.

#### Model 1 (Alpha)

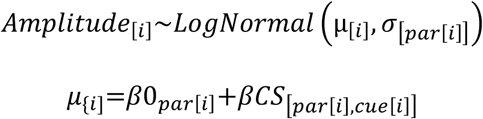

##### Multilevel priors

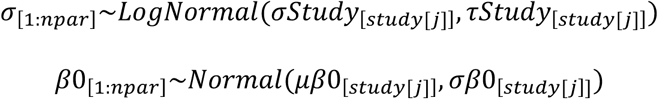

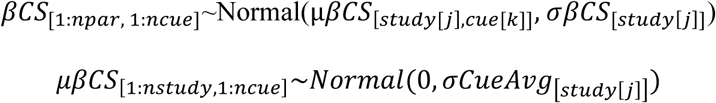

##### Priors (Weakly Informative)

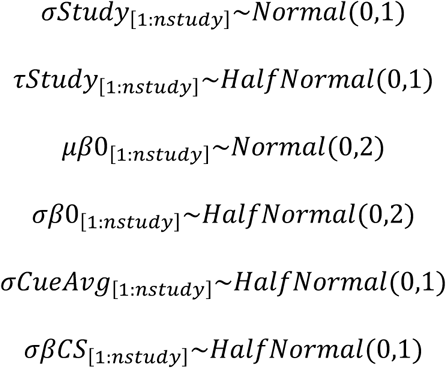

Model 2 for pupil amplitude was identical to Model 1 except that data was baseline-adjusted and each observation (i) was modeled based on a normal distribution with a mean, µ, and a standard deviation, σ.

##### Model 2 (Pupil)

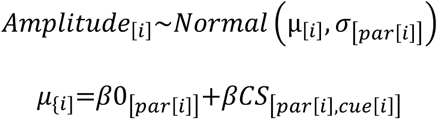

##### Multilevel Priors

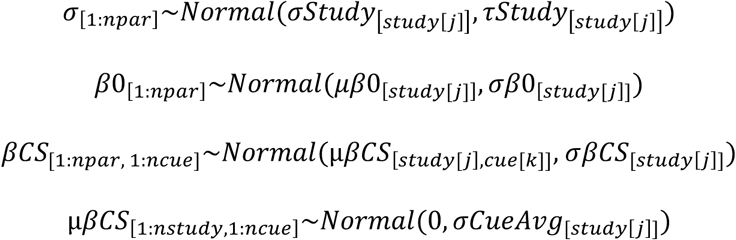

##### Priors (Weakly Informative)

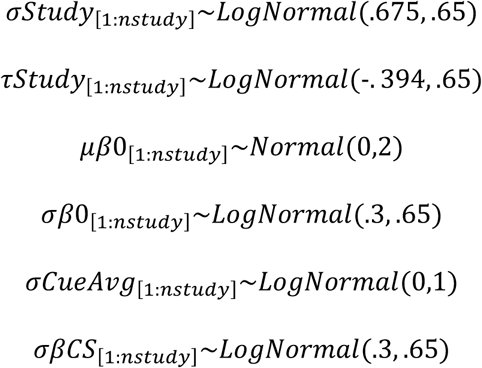

To understand how cue-evoked effects varied across experiments and physiological measures, we computed condition-and participant-level effect sizes posteriors from each model’s posterior draws. Because Bayesian models are generative, it is an accepted practice to create new posteriors from those estimated (McElreath, 2020). We define the effect size posteriors as the mean difference between CS+ vs CS-divided by the estimated single-trial error. This is similar to the effect size known as Cohen’s D, which is calculated as the mean difference between two groups over the pooled standard deviation (Cohen, 2013). The present approach is preferable to the standard Cohen’s D for a few reasons. First, it is more interpretable because it represents the effect size as a single-trial signal to noise ratio, rather than Cohen’s D, where group-level noise is influenced by the number trials averaged, which often differs in psychophysiological studies. Second, the effect size is estimated as a continuous posterior distribution, which conveys statistical uncertainty more effectively than a point-estimate like Cohen’s D. Finally, the effect size measures are more conservative and reliable, as both the mean differences and error were regularized through the multilevel structure. For group-level effect sizes, the average difference (µ*Cue*[CS+, Study[j]] – µ*Cue*[CS-,Study[j]]) was divided by the estimated error per study (σ*Study*[Study[j]]), whereas for each participant the subject level mean difference (βCS[Participant[k], CS+] - βCS[Participant[k], CS-]) via their estimated effects over their estimated error σ[Participant[k]]. Posteriors were summarized with the median and 95% credible interval as well as density visualizations (Figure 2 & 3).

## Results

Raw data for alpha and baseline-adjusted pupil diameter were extracted per trial and participant. The raw mean posteriors are shown in Figure 1. Both amplitude and pupil exhibited condition-dependent changes following stimulus onset as demonstrated by the posterior densities that differ for CS+ and CS-cues. For tone stimuli, there was evidence that the CS+ response exceeded that of CS-for pupil dilation with 99.8% (median = 0.08; [0.03, 0.14]) of the posterior contrast being above zero. Similarly, alpha amplitude showed 99.9% of CS+ draws exceeding the CS-draws (median = 0.07; [0.03, 0.11]). For face responses, there was stronger evidence that the CS+ exceeded CS-for both alpha reduction (98.5%; median = 0.06; [0.02, 0.09]) and pupil dilation (99.4%; median = 0.15; [0.06, 0.24]).

**Figure 1.**
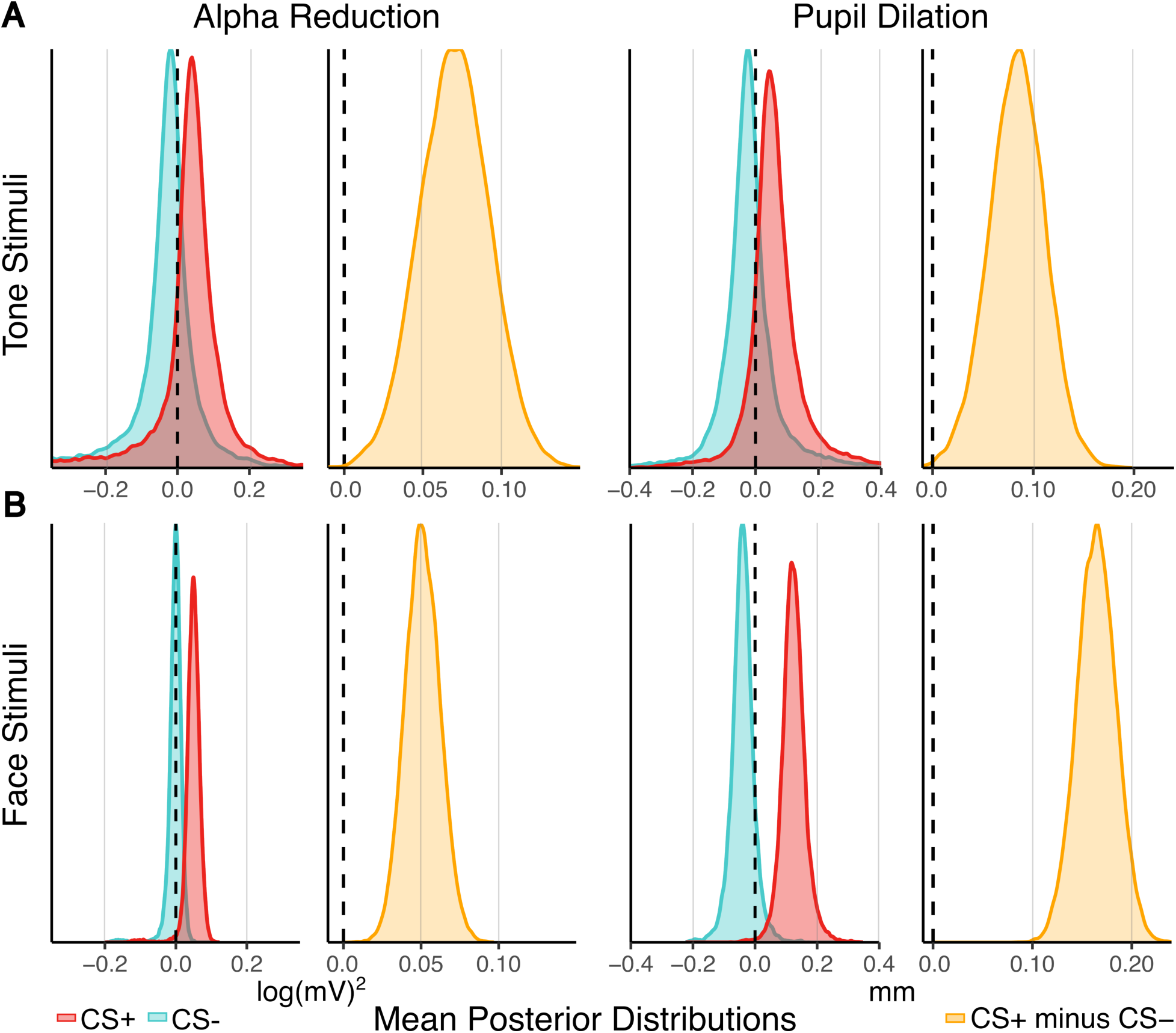
Posterior Means for Tone and Face Stimuli. *Note*. Posterior means for A) tone stimuli and B) face stimuli. Positive values indicate higher alpha reduction or pupil dilation response compared to baseline. Negative density values indicate less alpha reduction or pupil dilation response as compared to baseline.

Single trial effect size posteriors were estimated for alpha and pupil responses within each stimulus type (Figure 2, left panels). For tone stimuli, posterior median effect size for alpha was 0.12 [0.04, 0.19] and 0.14 [0.05, 0.24] for pupil. The posterior contrast between pupil and alpha for tone cues had a median of 0.02 [−0.10, 0.15], with 65% of the posterior density above zero suggesting similar effect sizes. For face stimuli, alpha effect size had a median of 0.13 [0.08, 0.19] and pupil had a median of 0.23 [0.05, 0.24]. The posterior contrast between pupil and alpha for face cues had a median of 0.07 [−0.01, 0.16] with 96% of the posterior mass above zero. Lastly, there was limited evidence that alpha or pupil differed across the two studies as the contrast between face study alpha and tone study alpha resulted in a median of 0.01 [−0.08, 0.12] while the contrast between face study pupil and tone study pupil resulted in a median of 0.06 [− 0.06, 0.18].

**Figure 2.**
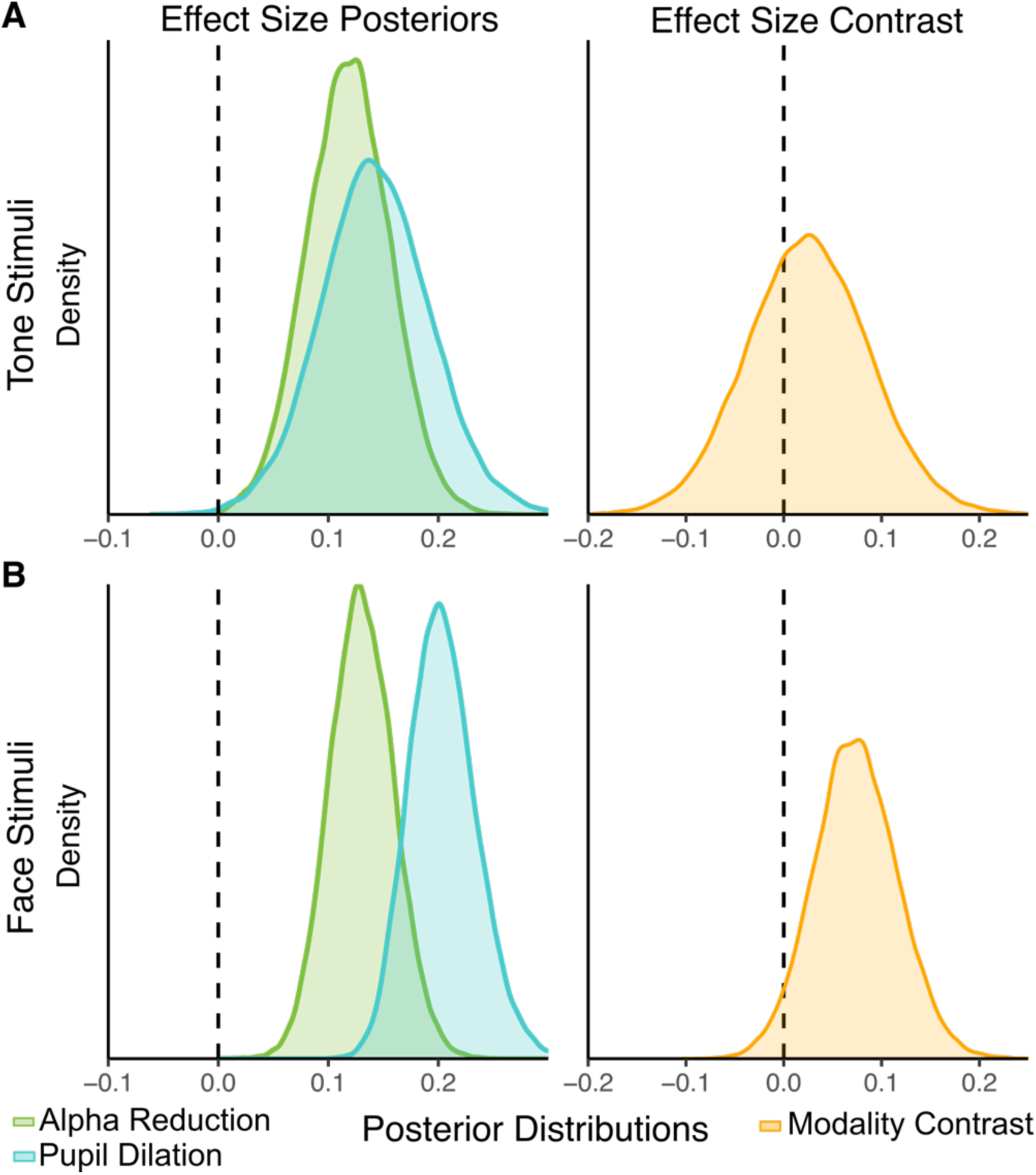
Group Level Single-Trial Effect Sizes Between CS+ and CS−. *Note.* Group-level single-trial effect sizes between CS+ and CS-for A) tone stimuli and B) face stimuli. The x-axis can be considered a signal-to-noise ratio between the mean difference and single-trial standard deviation. Results show similar effect sizes for tone stimuli mean posteriors demonstrated by the overlap in alpha and pupil distributions (A left panel) and posterior contrast straddling zero (A right panel). Face stimuli (B) show similar effect sizes to tone stimuli with posterior contrasts also overlapping with zero. There is mild evidence that pupil effect size to faces is larger than alpha with the posterior contrast showing 96% of effect sizes above zero (B right panel).

Effect size estimates were examined separately for each participant and stimulus modality (tone and face). Figure 3 illustrates the resulting per-participant distributions by measure, with distribution width reflecting uncertainty. Alpha (orange distributions) showed consistently smaller effect sizes, but also less variability, compared to pupil dilation (blue) distributions across both stimulus types. While the difference in effect size variability between the measures is apparent in Figure 3, this was quantified statistically by calculating posteriors of sample effect size variance. Per posterior draw, the variance function (var()) was used to find the effect size variance across the participants per measure and study resulting the desired posteriors. For the tone study, the effect size median variance for alpha was 0.001 [0.000, 0.004] and for pupil was 0.08 [0.04, 0.14] with the contrast greatly exceeding zero suggesting the effect sizes were more variable for the pupil measure (0.08 [0.04, 0.14]). This was also true for the face study in which alpha (0.001 [0.000, 0.002]) was less variable than the effect sizes for pupil (0.04 [0.02, 0.06]) with a contrast that exceeded zero (0.04 [0.02, 0.05]).

**Figure 3.**
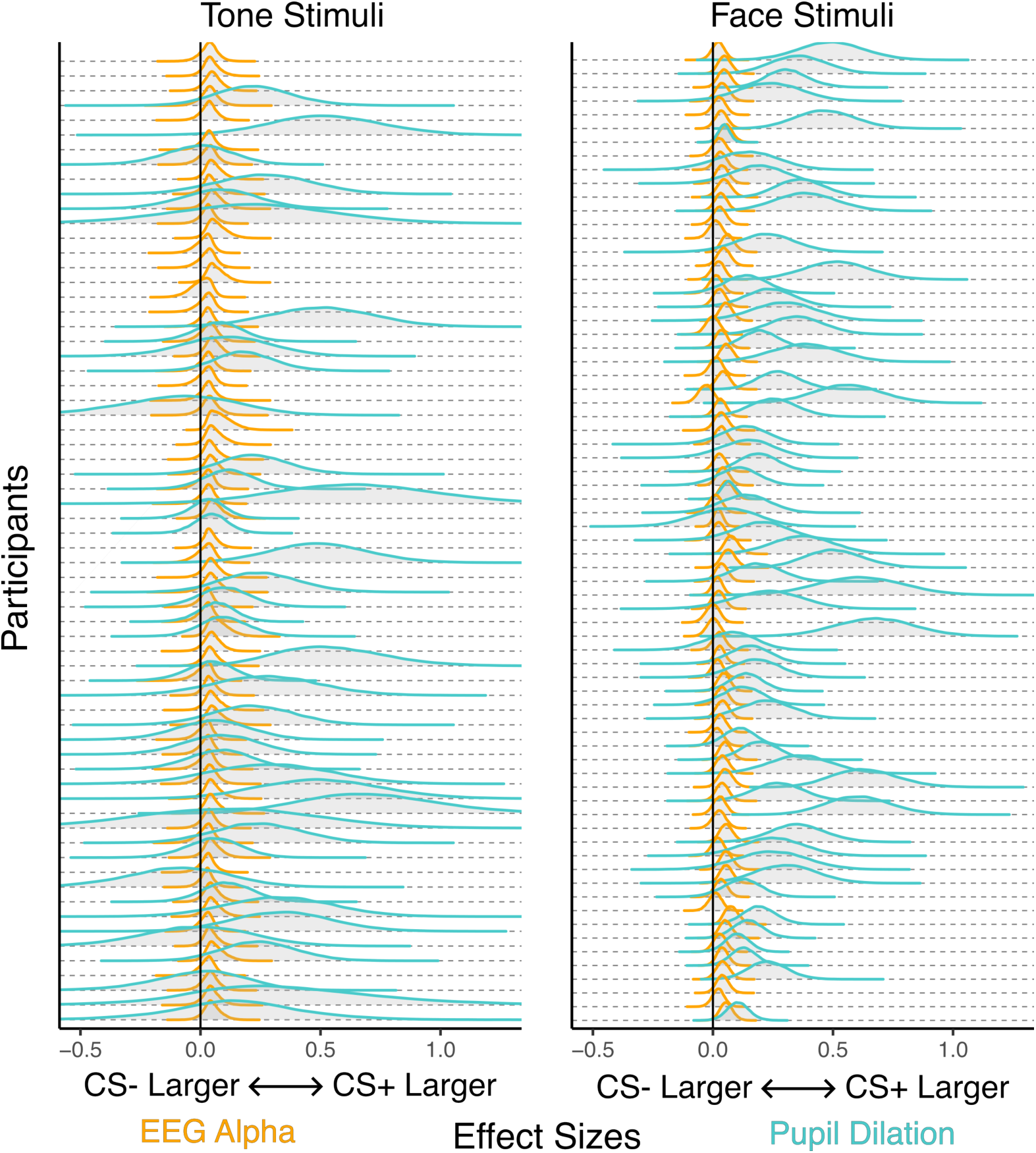
Per participant single-trial effect sizes by measure. *Note*. Per participant single-trial effect sizes by measure for tone stimulus in left panel and face stimuli in right panel. For tone and face stimuli, contrast is between effect sizes for CS+ and CS-over single-trial error (σ) per participant. Orange represents contrasts for alpha oscillations. Blue represents pupil dilation. Alpha shows more consistent effect sizes between participants. Pupil dilation shows more variation in effect sizes between participants, with less certainty in the effect size estimates.

## Discussion

The purpose of the current study was to characterize alpha and pupil variability within aversive conditioning at the single-participant level, across two simultaneous EEG and pupil conditioning studies with visual and auditory cues. Using Bayesian models, we quantified the extent to which conditioning effects differ within and between participants, considering alpha power and pupil diameter changes, and the two different conditioning paradigms.

First, we replicated previous group-level findings (Farkas et al., 2024; Pouliot et al., 2024) showing that aversive conditioning prompts selective changes in alpha power and pupil dilation. Posterior distributions of mean cue effects at the group-level showed the expected neural (EEG alpha power) and autonomic (pupil dilation) aversive conditioning responses; alpha power decreased while pupil dilation increased in the CS+ condition, compared to the CS-condition, in which the noxious US was never presented (Figure 1).

In order to compare the two physiological markers, differences between condition level posterior CS+ and CS-cues were divided by the estimated noise per-trial (Figure 2, left panels), providing an interpretable single-trial effect size posterior. For both tone and face stimuli, the posterior distributions across single-trial effect sizes for alpha power and pupil dilation were similar at the group level (Figure 2). The pupil effect size distribution (blue distribution) was slightly above the alpha power effect size distribution (green distribution), but the posterior contrast between the measures overlapped with zero, suggesting insufficient evidence that the measures were different (orange densities). The effect sizes posteriors were between .1 to .2 suggesting the mean difference was a tenth to a fifth as large as the standard deviation of random error per trial.

Next, the single-trial effect size posteriors were found per-participant to compare the variability between individuals (Figure 3). Evident in this visualization is that the similar group-level effects masked considerable differences at the individual-level. First, the conditioning effect on alpha power was found to differ very little between participants with nearly all participants having tight posteriors around an average effect size of 0.1. This finding is both novel and somewhat unexpected given the well-documented inter-individual variability in alpha power metrics (Gyurkovics et al., 2021a; Klimesch, 1999). For pupil diameter, while more participants had unusable data, there was also more variability between the conditioning effect sizes among people. For both paradigms, effect size posteriors were larger for pupil diameter contrasts. This was confirmed statistically by finding sample effect size variance posteriors and contrasting them between the measures. Previous literature suggests that this variability in pupil diameter may reflect individual differences in autonomic arousal and cognitive effort, as well as contextual influences that strongly affect the variability of sympathetic responses (Berntson et al., 1994; Steinhauer et al., 2022).

Understanding potential modality-specific effects is important for both theoretical and methodological reasons. While research in Pavlovian conditioning typically focuses on a single modality for CS+ cues, a complete neurophysiological account requires considering how different sensory systems, such as the visual and auditory systems, contribute to aversive learning. Bayesian hierarchical modeling provides a framework for comparing across experiments and modalities, enabling direct assessment of whether stimulus type influences discrimination between stimuli. This raises the question as to whether stimulus modalities index different components of an adaptive defensive response, rather than reflecting differences in arousal. From the perspective of motivational emotion theories (e.g., Frijda, 1988; Lang & Bradley, 2010), these measures may represent distinct action dispositions that support adaptive behavior. Pupil responses may be particularly informative in the context of visual stimuli such as faces, where rapid adjustments in visual sensitivity serve adaptive purposes. In contrast, selective suppression of the parietal alpha rhythm may reflect a more general attentional mechanism (Bacigalupo & Luck, 2022; Foxe & Snyder, 2011) that operates across modalities, supporting flexible engagement with both visual and auditory threat cues (Lang & Bradley, 2010).

Although evidence for modality effects on conditioned responses is limited, some research demonstrates that US modality shapes autonomic pupil responses (Finke et al., 2021). CS modality also impacts skin conductance responses (Sjouwerman et al., 2020; i.e., another physiological marker of arousal), and the cross-modal transfer of aversive conditioning responses between visual and auditory cues (Liu et al., 2025). These findings, as well as the results from our study, suggest that the nature of the US shapes the conditioned response, in line with traditional perspectives of Pavlovian conditioning (Pavlov, 1927; Rescorla & Wagner, 1972).

The current study found alpha band activity in EEG to show less variability between participants, although we did not find evidence that alpha and pupil differed at the group level. EEG signals are susceptible to noise from multiple sources, such as muscle artifacts, eye movements, and individual 1/f spectral slope differences, which may reduce the effect sizes for participant-level analyses (Gyurkovics et al., 2021b). While alpha effect sizes were more consistent across individuals, the magnitude of the effect in response to experimental manipulation is less robust compared to pupil response for a given participant.

There are several limitations to the present study. First, while multi-modal measurements provide additional information, their simultaneous use data may increase participant discomfort. Participant discomfort may manifest as increased movement resulting in greater artifact contamination, into the recordings. Second, the present studies did not utilize a chin rest, the use of which can negatively impact EEG data quality due to muscle artifacts. Conversely, using a chin rest can improve pupil data quality by reducing movement (Nyström et al., 2013). These methodological factors may explain some of the uncertainty in the pupil posterior distributions. Additionally, we assume homoscedasticity across trials – that is, error variance is presumed to remain constant over the course of the study. This assumption may be violated, especially in pupil measurements, where subtle shifts in participant posture or gaze can lead to transient loss of focus or misalignment with the eye-tracker, thereby introducing increased trial-dependent variability. However, baselining should correct for some of these experimental drifts in pupil measurements.

## Conclusion

Aversive conditioning research has traditionally emphasized group-level analyses, which provide an informative but incomplete account of psychophysiological effects. The current study utilizes Bayesian hierarchical modeling to compare alpha amplitude and pupil dilation at the individual level across two aversive conditioning experiments. At the individual level, alpha amplitude showed consistently small effect sizes across participants and studies, whereas pupil dilation exhibited more variable effect sizes. At the group level, there is marginal evidence that face stimuli produced a larger conditioning effect than tone cues suggesting pupil diameter may be more sensitive to direct sensory stimulation. Taken together, these findings imply that despite similar group-level effects, the choice of physiological measurement may depend on whether the goal of a given study is to identify consistent within-participant conditioning effects or to quantify the difference between individuals for clinical or translational research, in which robust descriptors of conditioning in each participant are sought. Future work could extend this approach to additional modalities and experimental contexts, helping to clarify how different physiological systems contribute to individual differences in aversive learning.

